# Increased female competition for males with enhanced foraging skills in Guinea baboons

**DOI:** 10.1101/2024.07.04.602040

**Authors:** William J. O’Hearn, Jörg Beckmann, Lorenzo von Fersen, Federica Dal Pesco, Roger Mundry, Stefanie Keupp, Ndiouga Diakhate, Carolin Niederbremer, Julia Fischer

## Abstract

Recognizing skillful group members is crucial for making optimal social choices. Whether and how nonhuman animals attribute skill to others is still debated. Using a lever-operated food box, we enhanced the foraging skill of a single male (*the specialist*) in one zoo housed and two wild groups of Guinea baboon (*Papio papio*). We measured group member’s behavioral responses before, during, and after our manipulation to reveal whether they focused on the outcome of the males’ actions or changed their assessment of his long-term value. During the manipulation, females in the specialist’s unit, but not the wider group, competed over access to the specialist - increasing their grooming of him tenfold and aggression near him four-fold. Both behaviors were predicted by the amount each female ate from the food box and returned to baseline within two weeks of its removal. This behavioral pattern supports an out-come-based assessment where females responded to male-provided benefits (utility) rather than attributing competence (value).

**Teaser:** Guinea baboon females monopolize males in relation to their current perceived utility.

## Introduction

Competence is a salient feature by which humans evaluate one another. Discerning others’ competence is relevant when choosing cooperation partners and when predicting competition outcomes (Fusaro et al., 2011; Hermes et al., 2016). In animals, competence is defined as the ability to effectively execute a collection of skills in suites of related tasks, i.e. foraging, mating, and fighting (Sih et al., 2019). For example, foraging competence comprises skills like choosing ripe fruiting trees, searching out birds’ nests, or opening hard crack nuts (Sih et al., 2019; Taborsky & Oliveira, 2012). Recognizing skills and attributing competences to group members serves an essential social function, as it allows individuals to watch for and interact with more skillful individuals to access the benefits they provide (Chapais, 2006; Ottoni et al., 2005; Whiten, 2018). Additionally, skill assessment in animals has been viewed as a precursor to human reputation and prestige, two traits that contribute to the formation and perpetuation of human cumulative culture (Chapais, 2015; Henrich & Gil-White, 2001; Whiten, 2018).

There is thus evidence that at least some animal taxa can recognize and respond to others’ skills. Some primate species pay more attention to proficient performers (*Pan troglodytes,* Kendal et al., 2018; *Cebus apella,* Ottoni et al., 2005), choose to cooperate with previously successful individuals (*Pan troglodytes,* Melis et al., 2006), or affiliate more often with individuals that can access unique foods (*Chlorocebus pygerythrus,* Fruteau et al., 2009; *Macaca fascicularis*, Stammbach, 1988). Some animals can also recognize human skillfulness. Female dogs (*Canis familiaris,* Chijiiwa et al., 2022) recognized which of the two human experimenters was more skilled at opening a food container. However, the cognitive mechanism by which skill attribution occurs remains unclear. Understanding how animals attribute skills can help us identify which features of social environments led to the development of sophisticated forms of competence attribution performed by humans (Henrich & Gil-White, 2001; Whiten, 2018).

We aim to distinguish between two core mechanisms. The first is an “outcome-based” process in which groupmates observe that a skilled individual’s actions can provide them with a beneficial outcome. The process is mediated by associative learning and increases the skilled individual’s perceived utility for as long as they can provide the desired outcome. Importantly, the outcome-based mechanism does not involve attributing competence to the individual. Instead, these individuals simply represent a source of greater utility (Hermes et al., 2015; Heyes, 2012). Following biological market theory, group members should compete for access to individuals with higher utility and possibly trade affiliation, such as grooming for access to the desired outcome (Noë & Hammerstein, 1994). Possible cases involving out-come-based mechanisms are experiments in which group members respond to an individual suddenly gaining sole access to a high-quality food source (Fruteau et al., 2009; Stammbach, 1988).

The alternative mechanism is “competence-based”, whereby group members observe a model individual and infer their behavior is indicative of skill in one or more domains – thereby attributing competence (Fusaro et al., 2011; Hermes et al., 2015). An inferred competence can be (i) narrow, creating only an expectation of skill at the same task – such as opening a palm nut – in the future (“behavior matching”); (ii) global, where a skill in one task is inferred as competence in all tasks (global evaluative thinking or “halo effects”); or (iii) flexible, where a skill in one domain – opening a palm nut in the domain ‘foraging’ – is understood as skill at related tasks in that domain, but not in others, e.g. fighting ability (“trait-based reasoning”; Fusaro et al., 2011). In all cases, a skilled individual is understood to have a higher value beyond its perceived utility in the current task as a result of their competence and should thus be preferred as a partner in future tasks relating to their competence (Brosseau-Liard & Birch, 2010; Cain et al., 1997; Fusaro et al., 2011; Hermes et al., 2015).

To shed further light on the cognitive processes by which animals evaluate the actions of others, we manipulated the skillfulness of a single individual and measured the behavioral responses of their groupmates to distinguish between outcome- and competence-based deci-sion-making. We conducted our manipulation in Guinea baboons (*Papio papio*) because their social environment may be conducive to partner evaluation based on competence. Guinea baboons live in nested multilevel societies (Patzelt et al., 2014), the base of which are stable ‘units’ comprised of a single reproductive male and one to several females with their young (Goffe et al., 2016; Pesco et al., 2022). Units are nested within ‘parties’, and parties are nested within ‘gangs’ (Fischer et al., 2017; Goffe et al., 2016). Within parties, males form enduring social bonds with other males (Dal Pesco et al., 2021), while male-female relationships are variable – some lasting years, others weeks (Goffe et al., 2016).

Guinea baboons are a promising system for investigating the role of skill in partner choice because they are generally relaxed and, excepting occasional conflicts, demonstrate extraordinary spatial tolerance (reviewed in Fischer et al., 2017). Thus, the payoff of a foraging skill acquired by one individual can be fed on by others with minimal conflict. Moreover, the society of Guinea baboons represents a novel social arena in which to test questions of social cognition where rank and kinship have only marginal effects on social relationships (Dal Pesco et al., 2021; Faraut et al., 2019; Kalbitzer et al., 2015). In the absence of strong rank-effects on social dynamics, we may be more likely to find individual partner preferences swayed by the outcome others provide, or their competences.

To distinguish between these two possible mechanisms, we artificially increased the foraging efficacy of one adult male Guinea baboon – the “specialist” (Stammbach, 1988) – in each of three Guinea baboon parties (one zoo housed, two wild), using a food box only the chosen individual could successfully operate. We measured the specialists’ social interactions before (baseline), during, and after a period of daily food box presentations to determine whether group members would alter their behavior in response to the specialist’s novel foraging skill. During the period of daily box presentations, we expected other group members to increase their affiliation with the specialist in response to either the benefit the specialist provided or in recognition of his competence. We were particularly interested in how group-mates would respond after the period of daily food box presentations. Persistent affiliation with the specialist in the weeks after the box was removed could reveal whether potential changes in the groupmates’ behavior towards the specialist were outcome-based or compe-tence-based. If group members were associating the specialist-box interactions with an outcome of desirable food, that is, the partner’s utility, then we expected affiliative behaviors to decay towards baseline, undergoing extinction once the box was no longer available. Alternatively, if group members were attributing a broader foraging competence to the specialist based on their success at manipulating the box (i.e. the inherent value of the specialist), we expected affiliative behaviors directed at the specialist to remain at elevated levels throughout the post-phase and possibly longer until group members would have had the opportunity to reassess the specialist’s competence.

## Methods

### Zoo site and study subjects

The first part of the study was conducted within behavioral enrichment procedures at Nuremberg Zoo (Tiergarten Nürnberg), Germany. The study group comprised 43 Guinea baboons, including 27 adults (10 males and 17 females), four sub-adults, and 16 immatures. Baboons in this group were experimentally naïve. Individuals were recognized by natural body markings and, for adult females, unique tattoos consisting of a letter and number on their left lower abdomen (see Supplemental Materials for details).

### Field site and study subjects

The fieldwork took place at the field station ‘Centre de Recherche de Primatologie (CRP) Simenti’ (13°01’34” N, 13°17’41” W) in the Niokolo-Koba National Park, Senegal (see Supplemental Materials for details). The study subjects belonged to two groups (“parties”) in different gangs. The home ranges of the parties covered, on average, 30.3 km^2^ of largely overlapping territories (Zinner et al., 2021). Party 5 comprised three adult males and eleven adult females, with one subadult male, one subadult female, and 24 juveniles arranged in four units. Party 6W comprised five adult males, six adult females, two subadult females, and 13 juveniles arranged in three units (Table S1). All subjects were fully habituated, naïve to experiments involving food, and individually identified via natural markings, body size and shape, and radio collars.

### Behavioral data collection

We collected data in the zoo from August to October 2021, which allowed us to assess the feasibility of our assay under more controlled conditions, and then in the field from January to August 2022 (see Supplemental Materials for details). Within each group, the study design comprised a training phase and three experimental phases: pre-, manipulation-, and post-phase. In the training phase, we trained several males in each group to operate the lever on the food box (zoo: three males, Party 5: one male, Party 6W: three males) and then selected one of the trained males to be our specialist for that group (see Supplemental Materials for details; Table S1). All specialists were mid-prime aged reproductive males with similar unit sizes (3-5 females). Once each specialist was chosen, we moved on to the three experimental phases.

In the pre- and post-phases, we only collected behavioral data. In the manipulation-phase, we collected behavioral data and presented the food box to the baboons each day. During all three phases of the experiment, we collected continuous focal follows (Altmann, 1974) of the specialist from 8:00 to 16:00 in the zoo, and from when we found the baboons in the morning, ∼7:00, to when we left the field ∼13:00. Importantly, we did not collect focal data during box presentations, but we video-recorded the full presentation to code feeding duration. During focal follows of the specialist, we recorded all grooming the specialist received and all instances in which another adult approached within 1 m of him. In the wild, we also recorded all conflicts instigated by a female member of the specialist’s unit that occurred within 5 m of the specialist ad-libitum. Unit-female aggressions were not initially part of the planned behavioral data collection. They were added after the experiment in the zoo, where we observed that females in the specialist’s unit appeared to be more aggressive during the manipulation-phase -starting more fights in and outside the unit. In all three groups we also recorded “party scans” in which every 20 minutes we took instantaneous scan sample of each member of the specialist’s party. In each scan we recorded the number and identity of all individuals with 1, 2, and 5 m of the subject at the time of the scan. Observational data were recorded with custom-made forms using the Pendragon 7.2 software (Pendragon Software Corporation, USA) running on cellular phones (Gigaset GX290, Gigaset, Germany).

### Zoo experimental procedure

Because we had no access to the enclosure’s interior, we installed a “food box” in the doorway to the old stable. The apparatus consisted of a single thick panel (0.6 m x 0.8 m x 0.015m) of macrolon with a single lever and two pipes, one short and one long, protruding from the side of the panel facing the enclosure (Fig. S2). An experimenter located behind the box could manually dispense peanuts down the pipes and trigger a speaker to play a tone after the lever was successfully pulled. The lever could be manually locked in place to prevent unwanted pulls.

In the zoo, each phase lasted ten working days. During the manipulation-phase, the box was presented twice daily at 10:00 and 15:00, two hours after feeding time, to maximize the baboon’s motivation. Presentations lasted either 15 minutes or the time it took for the specialist to pull the lever 20 times, which ever occurred first (trial length range: 10:15-15:00). Only the selected specialist was able to pull the lever successfully. If another individual approached the box, the experimenter locked the lever. After the specialist successfully pulled the lever, he was rewarded with three peanuts through the box’s short pipe. The surrounding individuals received twelve peanuts through the long pipe for each lever pull of the specialist (see Supplemental Materials for further details).

### Field experimental procedure

The food box (0.67 m x 0.43 m x 0.44 m) we used in the wild had an aluminum frame and 5mm clear plastic panels to allow the baboons to see the food (Figure 1; See Supple-mental Materials). The box’s lever could be locked with a remote control so that it could no longer be depressed.

**Figure 1.**
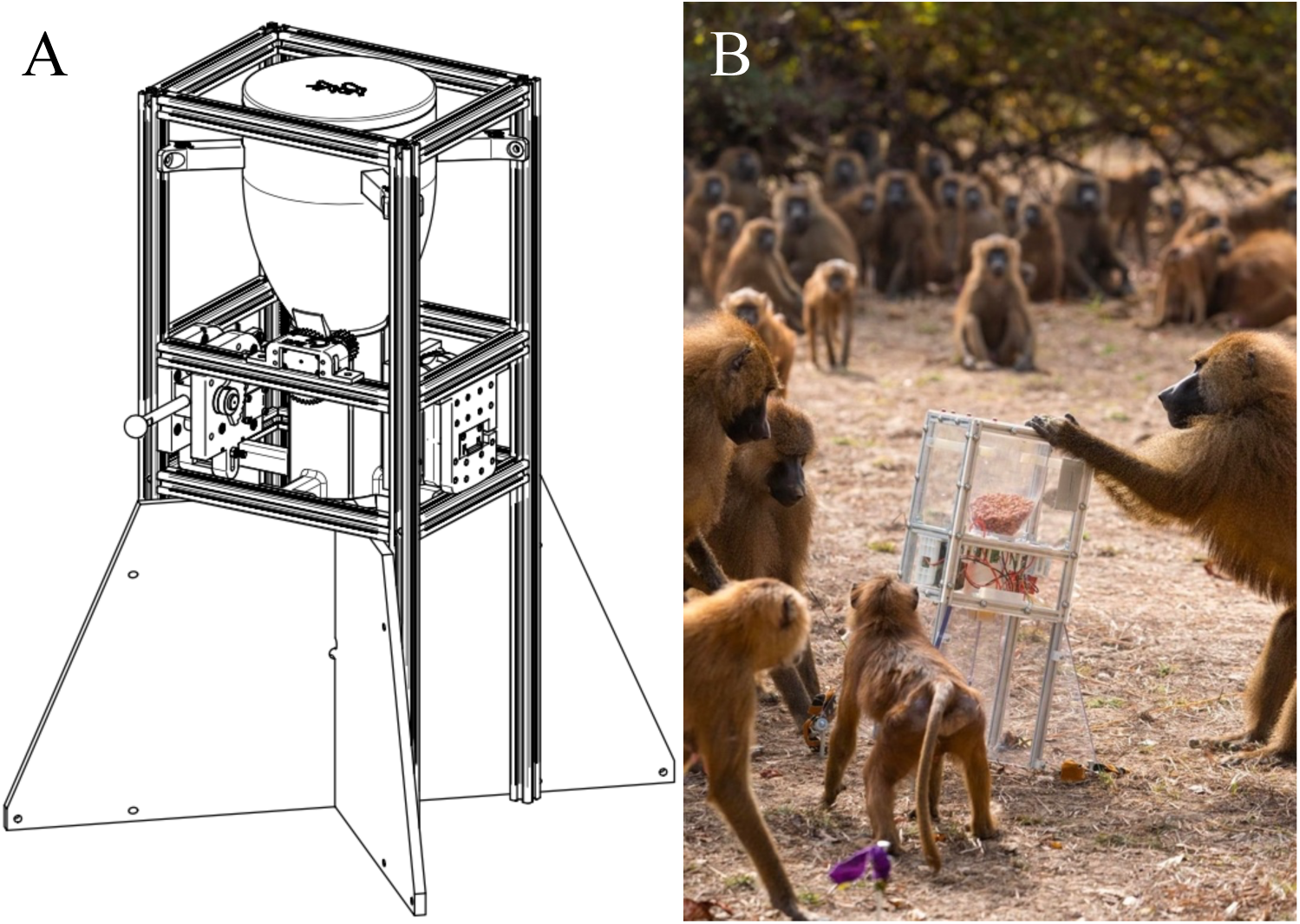
The food box, schematic by Louis Frank (left), used in presentations with the wild baboons (right), photo by Tessa Frank.

We increased the pre- and post-phases to twelve days each and the manipulation-phase to 15 days. The increased the phases lengths in the wild were because we could only present the box once per day, in the morning at the baboons’ sleeping site, and we needed a buffer against days when we might not find the animals (see Supplemental Materials for de-tails). Each time the lever was pulled, the box released ∼100 g of peanuts for the first nine pulls, which fell approximately evenly into the four ground regions separated by plastic di-viders. The tenth and final time the lever was pulled, it triggered a “bonanza”, releasing the remaining 1.3 kg of peanuts all at once, which formed a mound extending from the base of the box. The purpose of the ten pulls was to give other baboons more opportunities to see the specialist demonstrate his skill. The purpose of the “bonanza” was to create a widely distrib-uted, shareable food source that other baboons could access even after the specialist had left the box. A presentation ended when all the peanuts were gone (trial length range: 18:20-40:51) (see Supplementary Material for presentation details).

### Data analysis

We sought to measure if eating food from the specialist’s box altered the way individuals interacted with the specialist. To this end, we recorded the amount of food individuals ate from the box in each trial and interactions involving the specialist in the pre-, manipulation, and post-phases of the experiment. We were interested in three behaviors: 1) frequency of approaches to within 1 m of the specialist, 2) proportion of time spent grooming the specialist, 3) and frequency of aggressions initiated by unit females that began within 5 m of the specialist. Each response variable was modelled separately using Generalized Linear Mixed Models (GLMM; Baayen, 2008; McCullagh & Nelder, 1989). Each response variable was modelled separately using Generalized Linear Mixed Models (GLMM; (Baayen, 2008; McCullagh & Nelder, 1989)). The primary predictor variable in each model was an interaction of two fixed-effect terms. The first was the cumulative amount of time an individual spent eating food from the box each day (‘cumulative feeding time’). The second term was a variable capturing the number of days elapsed since the end of the manipulation-phase (‘post-day-number’). The post-day-number was zero for all days in the pre- and manipulation-phase, and then increased by one for each day in the post-phase (see Supplemental Materials for full model formulations).

We chose this set of predictor variables to model the predicted possible patterns in the response of the baboons, i.e. outcome-based vs competence-based. If group members’ inference was competence-based, indicated by levels of interactions with the specialist elevating in the manipulation-phase and remaining high in the post-phase (Figure 2 (a)), the resulting model coefficient for post-day-number would be close to zero, and the relevant term in the model that best explained the response would be solely cumulative feeding time. If group members’ inference was outcome-based and interactions with the specialist decreased in the post-phase, the coefficient for post-day-number would be negative, and cumulative feeding time would not strongly interact with post-day-number (Figure 2 (b)). Finally, we needed to account for the possibility that the decrease in the response in the post-phase would be expe-rience-dependent. For instance, benefitting more from the specialist’s skill could lead to a longer-lasting effect (response decreases more slowly; Figure 2 (c)) or the reverse (response decreases more rapidly; Figure 2 (d)). Such an effect can also be captured by including the interaction between post-day-number and cumulative feeding time in the model. Hence, the critical terms in the fixed effects part of the model were cumulative feeding time and its interaction with post-day-number. All terms included in an interaction were also included in the model.

**Figure 2.**
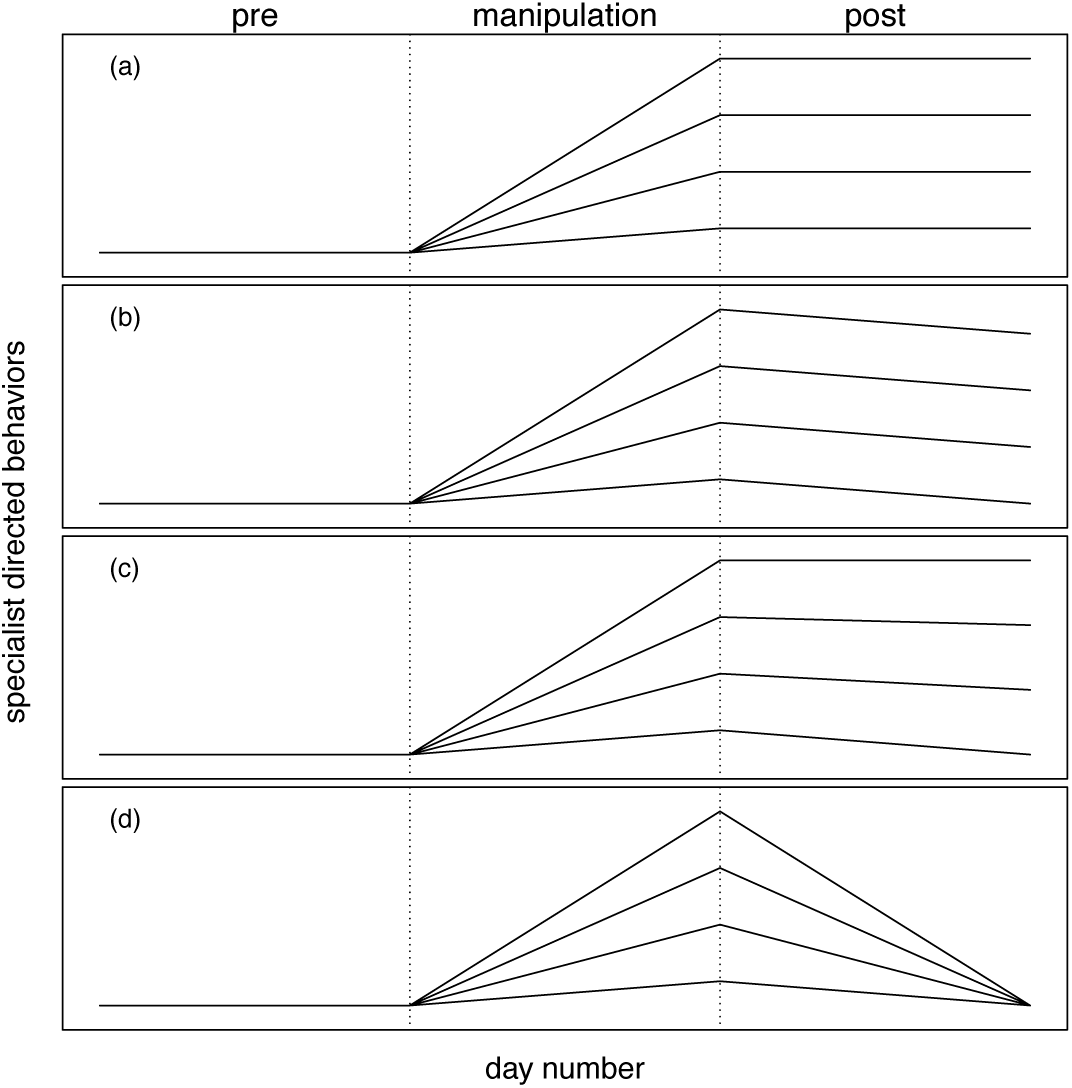
Illustration of possible patterns of specialist-directed interactions throughout the experiment. In the pre-phase, no systematic changes in interactions around the specialist are expected. During the manipulation-phase, the four fictitious individuals (each represented by one line) increasingly benefit from the specialist manipulating the food box, but to varying extents. Hence, one would expect the rate of behaviors directed towards the specialist to become increasingly common. In the post-phase, when the box is no longer available, there are several ways in which the interaction rates could change over time: (a) interaction rates could stay at the level that they had reached by the end of the manipulation-phase, (b) interaction rates could decrease at a similar pace for all subjects, (c) rates could decrease more slowly for subjects that interacted more in the manipulation-phase, or (d) rates could decrease more quickly for subjects that interacted more in the manipulation-phase to reach baseline levels at about the same time.

In the model where approach frequency was the response variable, we added the term ‘unit-member’ (yes/no) to the interaction between post-day-number and cumulative feeding, resulting in a three-way interaction between post-day-number, cumulative feeding, and unit-member. Unit-member was added to the model to account for the strong tendency of unit-members to approach their male. We initially had planned to include the term unit-member in all three models. However, grooming and aggression were only performed by members of the specialist’s unit, so it was unnecessary to include unit members in the other models.

In addition to our main predictors, we included precipitation as a fixed effect in all models. Precipitation was measured as mm of rain per day (Dal Pesco & Fischer, 2023). It was included to account for the tendency of the baboons, both zoo housed and wild, to form furry huddles whenever it rained -significantly altering interaction patterns. We also included subject and specialist IDs as random intercepts effects to account for repeated observations of the same partners and for differences between the three specialists. Lastly, to avoid the models being overconfident with regard to the precision of fixed effects estimates and to keep the type I error rate at the nominal level of 5%, we included random slopes of cumulative feeding time, post-day-number, their interaction, and precipitation within both grouping factors (Barr et al., 2013; Forstmeier & Schielzeth, 2011).

The model of approach frequency included all individuals present for at least one presentation of the box, which encompassed many individuals from other parties that had reduced access to the specialist due to parties splitting and merging over the course of the day. Thus, we included ‘duration present in group’ as an offset term (log-transformed, base *e*; (McCullagh & Nelder, 1989) to control for the time each individual was near the specialist and had the opportunity to approach. ‘Duration present in group’ was established by dividing the number of party scans in which a given individual was present by the number of party scans taken on that day and then multiplying the result by the duration of time for which the specialist was focal followed on that day. Similarly, some unit females may have had greater opportunity to instigate fights within 5 m of the specialist if they spent more time within these 5 m. To account for differing access to the specialist, we included as an offset term ‘duration present near specialist’ (log-transformed, base *e*), which was created by dividing the number of 5 m scans of the specialist in which a given female appeared by the number of specialist scans that day and multiplying the result by the focal duration of the specialist on that day. In this way, the measure of approaches and aggressions could be interpreted as the number of times an individual approached the specialist or started a fight near the specialist relative to their opportunity to perform either behavior.

The number of approaches toward the specialist was modeled as a count using a GLMM with Poisson error distributions using the R package lme4 (R version 4.2.0; R Core Team, 2022; lme4 version 1.1-21; Bates et al., 2015). Unit female aggressions were initially modeled as a count, but the initial Poisson model we attempted was overdispersed. We changed the model’s distribution to a negative binomial, which resolved the overdispersion issue. Grooming was modeled as the proportion of time that an individual spent grooming the specialist divided by the amount of time the specialist was observed using a beta distribution with glmmTMB (version 1.1.5, Brooks et al., 2017).

We inspected all quantitative predictors and responses for roughly symmetric distributions before fitting the models. We then divided cumulative feeding time and post-day-number by their maximum values, giving them a range from zero to one to ease model convergence and interpretation. We z-transformed precipitation to achieve the same end (Schielzeth, 2010). As an overall test of our main predictors and to avoid multiple testing, we conducted a full-null model comparison (Forstmeier & Schielzeth, 2011), whereby the null model lacked these cumulative feeding time and its interaction with post-day-number in the fixed effects part but was otherwise identical to the full model. The comparison was based on a likelihood ratio test (Dobson, 2002). After fitting the models, we checked the mean-variance assumption by checking whether the dispersion parameters deviated far from one (1.18, 0.94, 1.17). We determined Variance Inflation Factors (VIF) using the function vif of the package car (ver-sion 3.0-3; Fox & Weisberg, 2011) for a model lacking the interaction. Assessment of Variance Inflation Factors did not reveal any collinearity issues (maximum VIF: 1.197) (Quinn & Keough, 2002). We assessed model stability on the level of the estimated coefficients by excluding individual levels of the random effects (i.e., individual subjects and specialists) one at a time using a function provided by R.M. (Nieuwenhuis, 2012). The estimated coefficients did not vary substantially in the stability check, indicating the models were stable. We boot-strapped model estimates using the function bootMer of the packages lme4 and the function simulate of the package glmmTMB (N=1000 bootstraps).

## Results

We collected 554 focal hours from three specialists (one zoo housed, two wild). The sample included a total of 2481 approaches of 93 subjects toward three specialists. We found no significant effect of the interaction between cumulative feeding time, post-day-number, and unit member on the number of approaches individuals directed toward the specialist (full-null model comparison: χ^2^ = 5.21, df = 4, p = 0.266; Table 2). In other words, the frequency with which individuals approached the specialist was not related to the amount of food they ate from the food box.

Regarding grooming occurrence, 93 individuals were in the vicinity when the specialist operated the box, and 43 individuals ate food from the box. However, only 13 individuals groomed the specialist. In all three parties, the specialist was exclusively groomed by females within his unit. Contrary to our expectations, the specialist did not gain any new grooming partners as a result of our manipulation. Therefore, we fitted our grooming model using only data from females in the specialists’ units. The sample for the grooming model amounted to 466 observations of 13 females with three specialists. The interaction between cumulative feeding time and post-day-number had a significant effect on the duration of grooming directed toward the specialist (Estimate = −0.880, standard error = 0.383, *P* = 0.050; full-null model comparison: χ^2^ = 5.429, df = 2, *P* = 0.019). This result has two relevant implications. First, in the pre- and manipulation-phase, when post-day-number was zero, the amount an individual groomed the specialist increased as they ate more food from the box (Fig. 1). An increase of 60 minutes of peanut feeding (1 standard deviation) led to a nearly tenfold increase in grooming from 1.6% to 17.7% of the time the specialist was observed. Second, the negative coefficient for the interaction between post-day-number and cumulative feeding time (−0.880: Table 3) means that grooming decreased in the post-phase and that grooming decreased faster for females who fed more food or groomed more in the manipulation-phase. Put another way, the faster females increased their grooming in the manipulation-phase, the faster they decreased their grooming in the post-phase.

Interestingly, across all three groups, 42% (10.85/25.8 hours) of all feeding was done by individuals outside the unit, primarily by adult males from the specialist’s group. Adult males from the same group were among the most frequent feeders, eating more than any unit females in two of the three groups (Table 1). The males that ate the most were also frequently present in 5 m scans of the specialist (Table 1), reflecting their close social association with the specialist. Despite how much these males benefitted from the food reward, they did not change their behavior towards the specialist throughout the manipulation, as evidenced by their unchanged approach frequency and the complete absence of grooming.

The analysis of unit-female aggression contained only data from the two wild groups because we added this aspect only after our study in the zoo. The sample for this model includes 326 observations of ten subjects with two specialists. We found that the interaction between cumulative feeding time and post-day-number had a significant effect on the frequency of aggressions initiated by unit females (Estimate = −2.426, standard error = 1.106, *P* = 0. 020; full-null model comparison: χ^2^ = 7.610, df = 2, *P* = 0.022). Like the grooming model, this result has two relevant implications. First, aggressions and cumulative feeding increased together in the manipulation-phase. An increase of 67 minutes of feeding on peanuts (1 standard deviation) led to a nearly fourfold increase in aggressions instigated by female members of the specialist’s unit from 0.11 to 0.39 aggressions per hour. In other words, aggressions increased from one aggression every 10 hours to one every 2.5 hours. Sixty percent of aggressions (207/342) were directed at other females within the unit, 11.7% (40/342) at juveniles within the unit, and 27.8% (95/342) at individuals of both sexes beyond the unit. The second implication was that, like grooming, aggressions declined in the post-phase, and females that fed or aggressed more in the manipulation-phase decreased their frequency of aggressions faster in the post-phase. There was no significant effect of precipitation in any of the three models (Tables 2, 3, 4).

The stability checks of the grooming and aggression models showed that removing individuals from the analysis one by one did not change the results of the models but that removing specific individuals reduced the size of the effect that grooming or aggression had on cumulative feeding substantially (Table 3 & 4). Upon further inspection of the data, we found one to two females in each specialist’s unit that groomed, aggressed, and ate much more than other females (Table 1), indicating these females were primarily responsible for the effect measured in the models. These same effect-driving females also had the highest frequency of grooming and presence within 5 m scans of the specialist (∼20%), indicating they were the closest female associate or “favored” female of the specialist’s unit (Table 1). Thus, we found that the changes in grooming and aggressions were limited to the unit level. Within the unit, there were considerable disparities in the extent to which females modified their behavior toward the specialist.

As a post-hoc check for a general effect of provisioning on grooming rates (Gareta García et al., 2021), we analyzed behavioral data from 568 focal follows of 64 adults (37 females, 47 males) collected during the same period as the experiment, but outside box presentations. We found no correlation between the proportion of time individuals spent grooming others and the amount they fed from the box (r = 0.014). Furthermore, in both parties, the average proportion of time spent grooming others was no higher during the manipulation-phase than during the pre-or post-phases (Table 5).

**Figure 2.**
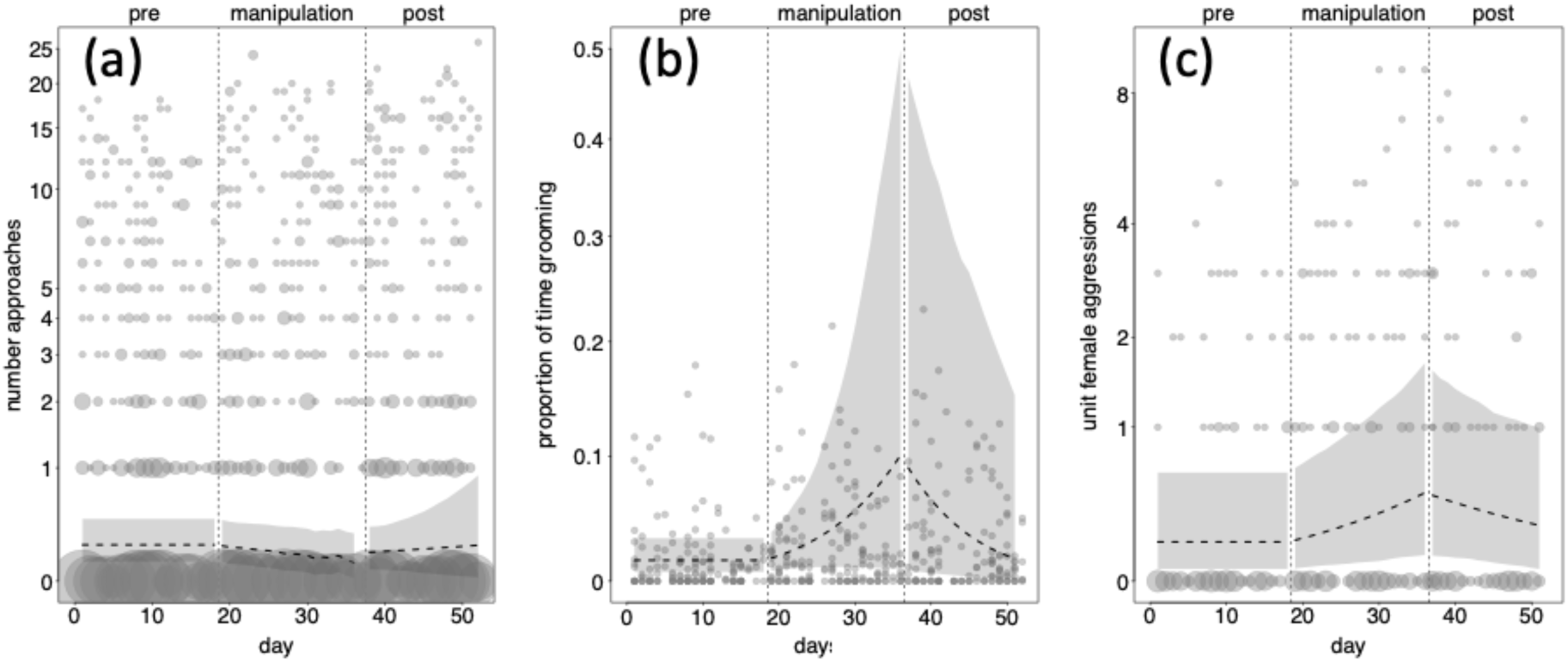
(a) Individual’s daily count of approaches towards the specialist, (b) proportion of individual’s daily time spent grooming the specialist divided by time the specialist was observed each day, (c) daily count of aggressions initiated by females in the specialist’s unit with 5 m of the specialist. Data represent the three experimental phases (pre, manipulation, post) separated by vertical dashed lines. The dashed horizontal dashed lines represent the fitted values with respect to cumulative feeding time and post-day-number (for the daily mean cumulative feeding time). The grey polygons around the dashed lines represent the 95% confidence interval from bootstraps around the fitted values for the mean cumulative feeding time in all three plots. The three y-axes are shown on the log scale to better differentiate the data points. The points show the raw data from wild and zoo housed groups (except aggressions, which we only collected in the wild), with the sample size per data point reflected by its area (range: 1 to 37).

**Table 1.**
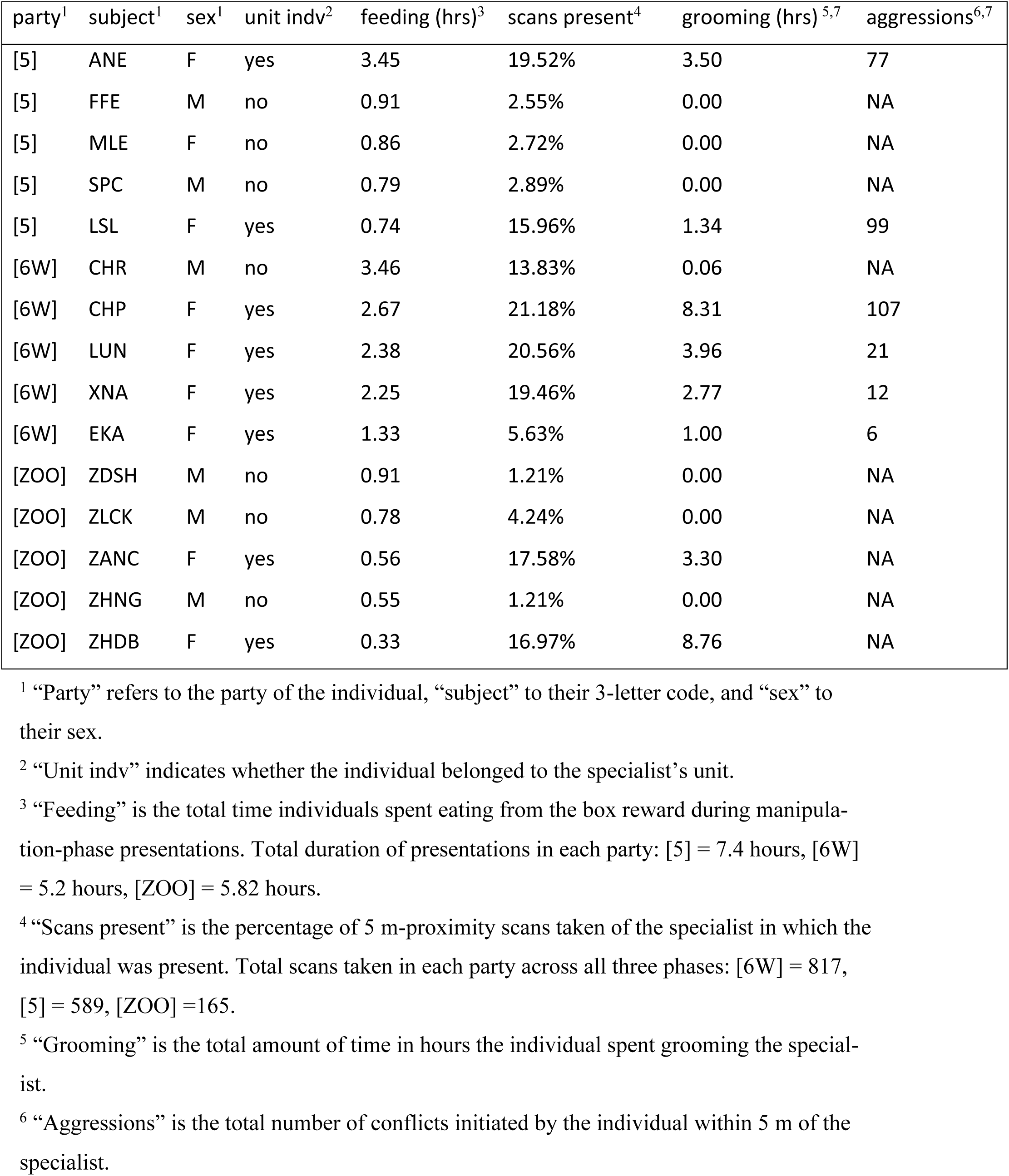
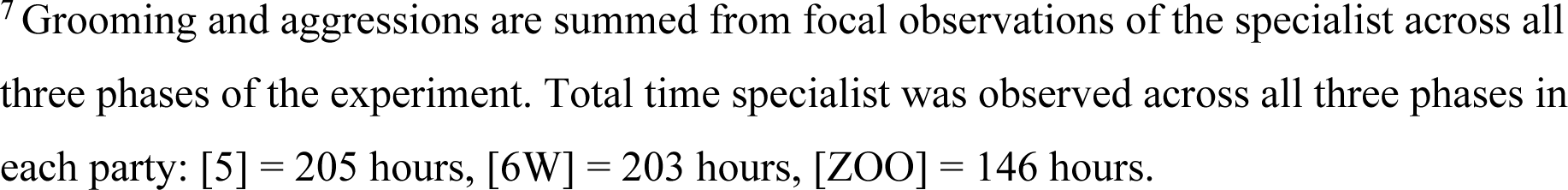
Summary statistics for the five individuals from each party, other than the specialist, who fed most frequently from the box.

| party<sup>1</sup> | subject<sup>1</sup> | sex<sup>1</sup> | unit indiv<sup>2</sup> | feeding (hrs)<sup>3</sup> | scans present<sup>4</sup> | grooming (hrs)<sup>5,7</sup> | aggressions<sup>6,7</sup> |
| --- | --- | --- | --- | --- | --- | --- | --- |
| [5] | ANE | F | yes | 3.45 | 19.52% | 3.50 | 77 |
| [5] | FFE | M | no | 0.91 | 2.55% | 0.00 | NA |
| [5] | MLE | F | no | 0.86 | 2.72% | 0.00 | NA |
| [5] | SPC | M | no | 0.79 | 2.89% | 0.00 | NA |
| [5] | LSL | F | yes | 0.74 | 15.96% | 1.34 | 99 |
| [6W] | CHR | M | no | 3.46 | 13.83% | 0.06 | NA |
| [6W] | CHP | F | yes | 2.67 | 21.18% | 8.31 | 107 |
| [6W] | LUN | F | yes | 2.38 | 20.56% | 3.96 | 21 |
| [6W] | XNA | F | yes | 2.25 | 19.46% | 2.77 | 12 |
| [6W] | EKA | F | yes | 1.33 | 5.63% | 1.00 | 6 |
| [ZOO] | ZDSH | M | no | 0.91 | 1.21% | 0.00 | NA |
| [ZOO] | ZLCK | M | no | 0.78 | 4.24% | 0.00 | NA |
| [ZOO] | ZANC | F | yes | 0.56 | 17.58% | 3.30 | NA |
| [ZOO] | ZHNG | M | no | 0.55 | 1.21% | 0.00 | NA |
| [ZOO] | ZHDB | F | yes | 0.33 | 16.97% | 8.76 | NA |

<sup>1</sup> “Party” refers to the party of the individual, “subject” to their 3-letter code, and “sex” to their sex.
<sup>2</sup> “Unit indiv” indicates whether the individual belonged to the specialist’s unit.
<sup>3</sup> “Feeding” is the total time individuals spent eating from the box reward during manipulation-phase presentations. Total duration of presentations in each party: [5] = 7.4 hours, [6W] = 5.2 hours, [ZOO] = 5.82 hours.
<sup>4</sup> “Scans present” is the percentage of 5 m-proximity scans taken of the specialist in which the individual was present. Total scans taken in each party across all three phases: [6W] = 817, [5] = 589, [ZOO] = 165.
<sup>5</sup> “Grooming” is the total amount of time in hours the individual spent grooming the specialist.
<sup>6</sup> “Aggressions” is the total number of conflicts initiated by the individual within 5 m of the specialist.
<sup>7</sup> Grooming and aggressions are summed from focal observations of the specialist across all three phases of the experiment. Total time specialist was observed across all three phases in each party: [5] = 205 hours, [6W] = 203 hours, [ZOO] = 146 hours.

**Table 2.** Output for the model with approach frequency as the response.

| Term | Estimate ± SE | Lower CI | Upper CI | $\chi^2$ | df | P | Min | Max |
| --- | --- | --- | --- | --- | --- | --- | --- | --- |
| intercept | -2.591 ± 0.332 | -3.346 | -2.161 | | | | -3.003 | -2.556 |
| cumulative feeding time<sup>1</sup> | -6.228 ± 3.578 | -19.688 | 0.358 | | | | -10.977 | 0.393 |
| post-day-number<sup>1</sup> | 0.010 ± 0.287 | -0.588 | 0.276 | | | | -0.271 | 0.259 |
| unit-member | 3.927 ± 0.444 | 2.900 | 4.686 | | | | 3.334 | 4.168 |
| precipitation<sup>2</sup> | -0.045 ± 0.987 | -0.078 | 0.112 | 0.605 | 1 | 0.608 | -0.134 | 0.057 |
| cumulative feeding time:post-day-number:unit-member | -5.852 ± 4.200 | 16.623 | 5.735 | 1.331 | 1 | 0.164 | -8.315 | 0.906 |

<sup>1</sup> cumulative feeding and post-day-number were divided by their maximum value to make them range from 0 to 1. The maximum values were cumulative feeding time = 12459.8, post-day-number = 15
<sup>2</sup> precipitation was z-transformed to a mean of zero and a standard deviation of one; the origi-nal mean was 2.75, and the standard deviation was 7.40

**Table 3.** Output for the model with the proportion of time grooming the specialist as the response.

| Term | Estimate ± SE | Lower CI | Upper CI | $\chi^2$ | df | P | Min | Max |
| --- | --- | --- | --- | --- | --- | --- | --- | --- |
| intercept | -4.115 ± 0.381 | -4.924 | -3.369 | | | | -4.630 | -3.900 |
| cumulative feeding time<sup>1</sup> | 2.579 ± 1.730 | -0.949 | 6.066 | | | | 0.839 | 4.014 |
| post-day-number<sup>1</sup> | 0.007 ± 0.169 | -0.340 | 0.342 | | | | -0.142 | 0.259 |
| precipitation<sup>2</sup> | -0.032 ± 0.062 | -0.153 | 0.083 | 0.284 | 1 | 0.594 | -0.335 | 0.006 |
| cumulative feeding<br>time: post-day-number | -0.880 ± 0.383 | -1.477 | -0.338 | 3.856 | 1 | 0.050 | -1.882 | -0.395 |

<sup>1</sup> cumulative feeding and post-day-number were divided by their maximum value to make them range from 0 to 1. The maximum values were cumulative feeding time = 12459.8, post-day-number = 15
<sup>2</sup> precipitation was z-transformed to a mean of zero and a standard deviation of one; the origi-nal mean was 2.75, and the standard deviation was 7.40

**Table 4.** Output for the model with aggression frequency as the response.

| Term | Estimate ± SE | Lower<br>CI | Upper<br>CI | $\chi^2$ | df | P | Min | Max |
| --- | --- | --- | --- | --- | --- | --- | --- | --- |
| intercept | -2.205 ± 0.651 | -3.635 | -0.999 |  |  |  | -2.535 | -1.765 |
| cumulative feeding<br>time<sup>1</sup> | 1.277 ± 0.355 | 0.286 | 2.206 |  |  |  | 0.859 | 1.638 |
| post-day-number<sup>1</sup> | 1.024 ± 0.573 | -0.607 | 2.106 |  |  |  | 0.558 | 1.301 |
| precipitation<sup>2</sup> | -0.021 ± 0.104 | -0.245 | 0.178 | 0.187 | 1 | 0.655 | -0.376 | 0.036 |
| cumulative feeding<br>time:post-day-number | -2.426 ± 1.106 | -4.982 | -0.080 | 5.407 | 1 | 0.020 | -2.788 | -0.449 |

them range from 0 to 1. The maximum values were: cumulative feeding time = 12459.8;
post-day-number = 15
<sup>2</sup> precipitation was z-transformed to a mean of zero and a standard deviation of one; the origi-nal mean was 2.75, and the standard deviation was 7.40

**Table 5.**
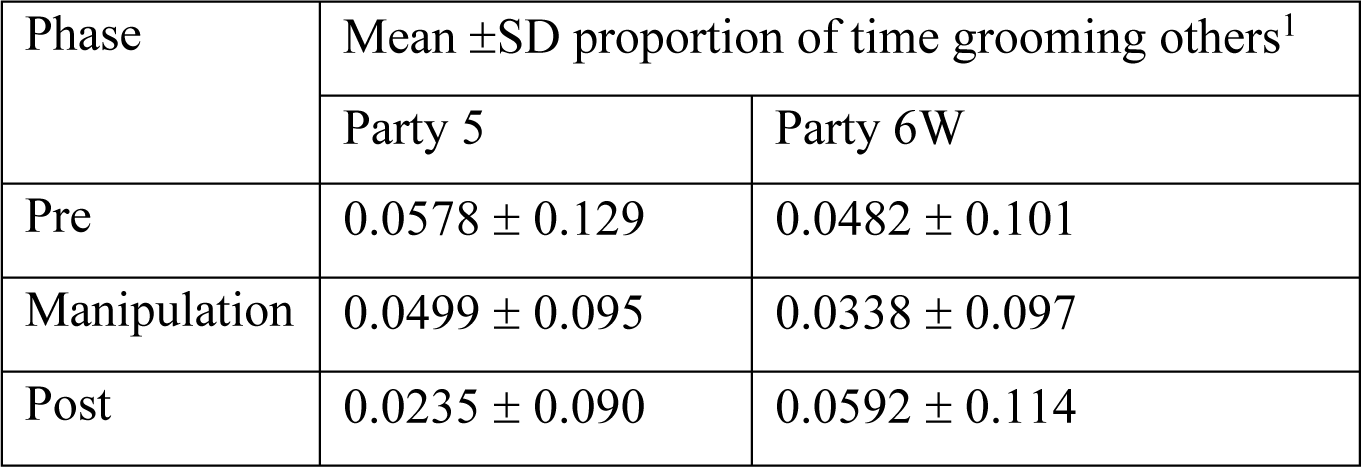
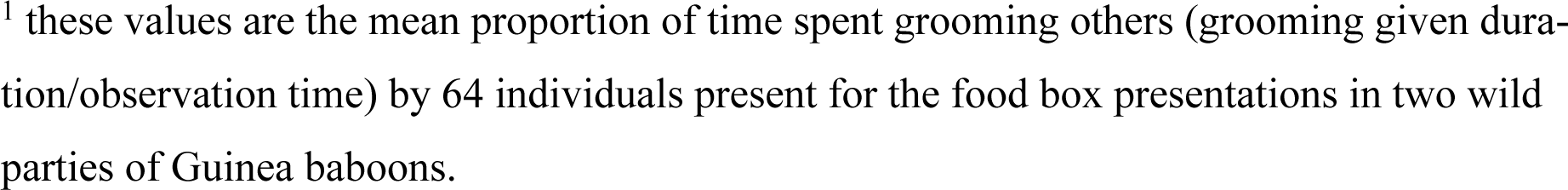
Party-wide grooming averages.

| Phase | Mean ±SD proportion of time grooming others<sup>1</sup> |  |
| --- | --- | --- |
|  | Party 5 | Party 6W |
| Pre | 0.0578 ± 0.129 | 0.0482 ± 0.101 |
| Manipulation | 0.0499 ± 0.095 | 0.0338 ± 0.097 |
| Post | 0.0235 ± 0.090 | 0.0592 ± 0.114 |

<sup>1</sup> these values are the mean proportion of time spent grooming others (grooming given duration/observation time) by 64 individuals present for the food box presentations in two wild parties of Guinea baboons.

## Discussion

In all three parties, we observed a dramatic change in behavior by the females within the specialists’ units. Unit females sharply increased the amount of time they spent grooming the specialist and the frequency with which they initiated aggressions near him. Escalations in both behaviors corresponded with the cumulative amount of food each female ate from the food box, which the specialist alone could open, and then reduced to baseline values after the box was taken away. This pattern is consistent with our predictions for an outcome-based process wherein of the unit-females’ respond to the temporarily elevated utility of their male. In a post-hoc test, we found that the amount group members groomed individuals other than the specialist was not correlated with the amount they fed from the box. Thus, the changes in grooming directed toward the specialist were not simply the result of provisioned individuals having more time to spend grooming but were instead related to the perceived increase in the utility of the specialist. To our surprise, there was no widespread change in affiliation toward specialists, i.e., at the group level, as observed in previous “specialist” manipulations (Fruteau et al., 2009; Stammbach, 1988). Individuals did not approach the specialist more often, nor did the specialist gain new grooming partners due to his access to the novel food source.

Interestingly, despite males from the specialist’s group feeding prodigiously, there was no apparent change in the frequency with which they approached or interacted with the specialist. Hence, in this experiment, we saw highly differentiated responses between males and females, which correspond to the social stratification of this multilevel society -wherein male allies already “earned their place at the table”. In contrast, associated females fiercely competed over their males who had gained in utility.

### Female competition within the unit

At the unit level, we saw changes in behavior indicative of increased competition between unit females over the specialist. In response to his novel foraging skill, females groomed the specialist more to secure their position with the male [attitudinal reciprocity (Brosnan & Waal, 2002; De Waal, 2000; Schino & Aureli, 2010)] and aggressed others nearby to prevent them from doing the same [see grooming intervention (Massen et al., 2014; Mielke et al., 2017, 2020; Schneider & Krueger, 2012)]. Thus, the increase in grooming is not tied to getting more food per se, but firstly to establishing a top position with the male (so long as he provided resources), and then, as a result of that position, gaining better access to the food he controls. In this sense, the females’ attitudes towards the male changed as he became a more desirable resource worth aggressively defending against other females – both inside and outside the unit. Notably, the grooming and aggressive behavior of unit females did not occur during presentations of the box and instead took place over the course of the day. Thus, the behaviors were not a tit-for-tat exchange of grooming for tolerance at the food site. Instead, they represented a broader response to the change in their male’s inherent utility.

In societies based on one-male-units such as geladas *Theropithecus gelada* (Crook, 1966), *Gorilla gorilla gorilla s* (Robbins et al., 2004), *Rhinopithecus roxellana* (Qi et al., 2009), *Equus caballus* (Schneider & Krueger, 2012), and *Papio papio* and *Papio hamadryas* (Fischer et al., 2017; Kummer, 1968), reproductive males confer a range of benefits on females in their units. For instance, social proximity to the reproductive male enhances protection from predators (Dunbar, 1983; Sigg, 1980) and grants priority access to mating opportunities (Harcourt, 1979; Qi et al., 2011), general food sources (Sigg, 1980), rare or desirable food sources (Goffe & Fischer, 2016), and paternal care (Harcourt, 1979). The reproductive male is also the most valued coalition partner in the unit – greatly enhancing females’ abilities to win conflicts against opponents both in and outside the unit (Dunbar, 1983; Fischer et al., 2017; Harcourt, 1979; Kummer, 1968). However, these benefits are often not distributed equally, with females possessing the strongest ties to the male reaping most of the benefits (Dunbar, 1983; Harcourt, 1979; Kummer, 1968; Qi et al., 2011). In line with these previous findings, we conclude that female competition over their male is likely the primary force explaining the responses to the males’ experimentally enhanced foraging skills.

### Male tolerance at the group level

We saw no effect of our manipulation on the behavior of individuals outside the specialist’s unit. Despite their unchanged behavior, males from the specialist’s group were among those individuals that benefited most from his lever-pulling skill. The ability of male group members to tolerantly feed shoulder to shoulder from food that unit females were competing over suggests radically different social dynamics for the in- and out-group members, i.e., at the level of the unit vs. the party. The strong male-male bonds that link Guinea baboon units together at the party level (Dal Pesco et al., 2021; Patzelt et al., 2014) also permit males tolerance and access to rare high-quality food sources. If this is the case, males do not need to monitor the foraging skill of other males when they can simply monitor the presence of food and then fed from it regardless of its possessor (Table S2).

### Cognition of skill

The primary goal of our study was to investigate the cognitive processes underpinning the responses to a group member’s enhanced foraging skill. To this end, assessing group members’ behavior in the post-phase was definitive. Crucially, when the period of box presentations ended, females reduced their affiliation and aggressions, returning to baseline frequencies. The observed pattern indicates an outcome-based response where the partner’s utility increased as a result of others benefitting from him operating the box. Once the box was removed and the specialist could no longer use his skill to provide food, the association between him and the outcome began to decay and, without further reinforcement, reached extinction (Rescorla, 1988). Moreover, in our study, the more a female within the specialist’s unit benefitted from the outcome of his competence, the more she altered her behavior around him (i.e. grooming, aggression). This finding suggests that the individual changes in behavior were based on the extent of positive outcomes they experienced. This process differs from competence-based inference, such as behavior matching, halo effect, and trait reasoning, where indirect experience can yield the same assessment of competence as direct experience, despite never receiving any benefit from another’s competence (Fusaro et al., 2011; Hermes et al., 2016).

Competence-based trait attribution is a basic tenant of human social interactions, with children as young as four combining global evaluative judgements with trait-like inferences (Cain et al., 1997; Hermes et al., 2015). However, while there is evidence that nonhuman animals can recognize and respond to the skills of conspecifics (Chijiiwa et al., 2022; Fruteau et al., 2009; Ottoni et al., 2005; Stammbach, 1988), to our knowledge, only one other study has examined how those skills were attributed. Keupp and Herrmann (2024) found that captive chimpanzees use a simple competence-based process (behavior matching) to evaluate the skill of conspecifics in choosing a collaboration partner. Our results contribute to our emerging understanding of skill attribution in nonhuman primates by demonstrating that Guinea baboons use an outcome-based process in response to a novel foraging skill in a groupmate. Though the contexts of the foraging tasks differed between the two studies, our results imply a gap between the type of inference baboons and chimpanzees used to assess skills in the foraging domain (Keupp & Herrmann, 2024).

In summary, we conclude that female Guinea baboons attribute the change in their male’s foraging skill using an outcome-rather than a competence-based process. Our results highlight how individuals’ social strategies are shaped by the utility of available social partners and the organization of the society in which they are embedded. Additionally, we observed the same responses to our manipulation in both a zoo housed and wild population of Guinea baboons, indicating the mechanism underlying the response is generalizable between population and environment. Broader comparative studies across multiple taxa and different cognitive domains will be needed to shed light on the evolution of competence attribution and identify which social contexts promoted its emergence.

### Ethics Declaration

This research approved by the Senegalese Direction des Parcs Nationaux (Protocol d’Accord entre DPN and DPZ signed 22.4.2019) and conforms with the guidelines for the ethical treatment of nonhuman animals set down by the Association for the Study of Animal Behaviour (ASAB Ethical Committee/ABS Animal Care Committee, 2023).

## Acknowledgments

We would like to thank the Direction des Parcs Nationaux (DNP) and the Ministère de l’Environnement et de la Protection de la Nature (MEPN) du Sénégal for approval to conduct this study in the Parc National du Niokolo-Koba (PNNK). We particularly thank the former conservateur of the park Jacque Gomis for his cooperation and logistical support; all the staff and field assistants of the CRP Simenti, in particular Irene Gutiérrez Díez, Lisa Orndorf, Marc Mönich, and Rachel Sassoon for their help with the data collection; and all the staff of the Tiergarten Nürnberg team, Dagmar Fröhlich, Ramona Such, and the zoo technicians for their support, as well as Hugo Matthes for his tireless efforts. This research was funded by the Deutsche Forschungsgemeinschaft (DFG, German Research Foundation) – Project-ID 454648639 SFB 1528 “Cognition of Interaction “and Project-ID 254142454/GRK 2070 “Understanding Social Relationships”. Support by the Leibniz ScienceCampus “Primate Cognition” (Audacity Fund AF2020_03), and by the DAAD “One-Year Grants for Doctoral Candidates 2020/21 (57507870)” are gratefully acknowledged.

## Author contributions

W.O. and J.F developed the concept and design of the study; W.O. designed and built the boxes; J.B. and L.v.F provided access to zoo housed baboons. W.O., N.D., and C.N. conducted the fieldwork; F.D.P. assisted with data preparation; W.O. and R.M. designed and conducted the analysis; F.D.P., J.B. and L.v.F provided conceptual input; W.O. drafted the manuscript; W.O., J.B., L.v.F, F.D.P, R.M., S.K., N.D., C.N., and J.F. reviewed the manuscript and provided final approval of the submitted version. W.O. and J.F. acquired the funding.

## Data availability statement

There is no dataset and code associated with this manuscript can be found at https://osf.io/2fnvd/?view_only=f42cdd8465e54e0686abdb0978fc109a.

