## Supplemental Materials for "Increased female competition for males with enhanced foraging skills in Guinea baboons"

**This PDF file includes:**

Supplementary Text
Figs. S1 to S3
Tables S1 to S2

**Other Supplementary Materials for this manuscript include the following:**

Movies S1 to S3

**Supplementary Text**

Zoo site and study subjects

The age and sex class of all individuals in the population were determined using the categories from the CRP Simenti Work Manual (Dal Pesco & Fischer, 2023). Individuals over six years of age were considered adults or sub-adults, named, assigned a unique four-letter code, and were subject to data collection.

The baboons' enclosure consisted of an outdoor area (ca. 310 $m^2$), the "new stable" (ca. 94 $m^2$), and the "old stable" (ca. 18 $m^2$) (Fig. S1). The old stable was not accessible to the animals but contained a hatch through which we presented the food box.

Field site and study subjects

The field site lies next to the Gambia River, where multiple seasonal wetlands (Mares) occur in depressions alongside the river. The prevailing vegetation types are dry forests and various savannah types, including savannah woodlands, tree/shrub savannahs, and grass savannahs (Zinner et al., 2021). The climate is highly seasonal, with a dry season from November until May and a rainy season from June until October (Fischer et al., 2017).

Behavioral data collection

Before the start of the experiment, all experimenters were trained on the ethogram and data collection methods used at the Simenti field site in Senegal (Dal Pesco & Fischer, 2023). Data collection began after all experimenters passed an ID and inter-observer reliability test (see Supplemental Materials for details).

Zoo experimental procedure

Unfortunately, the specialist broke the mount for the long pipe during the third trial. Instead of replacing the mount, we placed the same long pipe at the edge of the enclosure, positioned 5 m above and directly behind the specialist for all subsequent trials to shower peanuts over a wide area (~2m2) behind the specialist at every pull. Presentations of the box were recorded with two GoPro Hero 8 video cameras (GoPro, California, United States), one placed on the inner side of the box facing outward and the other placed above the enclosure facing the area in front of the box, resulting in an overhead view.

Field experimental procedure

At the start of the presentations, the food box was staked to the ground, and three colorful marker stakes were placed in a semi-circle 2 m and 5 m from the box (See image of set up Fig. S3). The flags aided in estimating distances during subsequent video coding. The box contained ~2.2 kg of peanuts without their shells because the dispensing device was incompatible with the peanuts in their shells used in the zoo. As a result, lever pulls yielded more edible material and thus wild baboons got the same amount of food as captive baboons with only one presentation per day and only 10 lever pulls.

Video coding

Two researchers conducted presentations. One researcher operated a stationary GoPro Hero 8 on a tripod, which we placed facing the lever side of the food box. This researcher also operated the food box remote control so they could lock or unlock the lever to ensure only the specialist pulled the lever. The second researcher stood opposite the stationary camera and used a handheld Panasonic HC-X909 video camera (Panasonic Corporation, Kadoma, Japan) to record individuals on the back side of the box and the wider audience behind the stationary camera. The second researcher also verbally identified all baboons within sight for later use in video coding

Presentation videos were coded by placing the two recordings (zoo: overhead and box view; field: stationary and handheld) side by side and synchronized in time within a single video using VSDC Free Video Editor 7.1 (see Example Trials). Two observers coded the amount of time each individual spent actively feeding on peanuts from the food box to the nearest third of a second using Solomon Coder version beta 19.08.02. To ensure interobserver reliability, both observers coded a selection of six videos in both the zoo and field settings. There was a high degree of agreement (91%) in the assessment of feeding duration between observers (see Supplemental Materials for details).

Inter-observer reliability test

Once ID training came to an end and researchers were capable of reliably identifying the baboons and collecting data, we checked observer reliability by conducting parallel observations of at least ten 20-minute focal protocols with another researcher. We then compared the protocols of the two observers and checked whether they had the same entries. We established the percentage agreement between the two observers by dividing the number of lines in which their entries agreed (A) by the number of the lines in which the observers disagreed or were missing (D) (e.g., $X = A/(A+D)$; see Bateson & Martin, 2012). Newly trained researchers were ready to focal on their own if the percentage agreement was above 80% (= 0.80).

We used the same process for video coding, comparing each line (1 for each third of a second) for each coded individual where the two coders marked the animals as feeding or not feeding.

Training phase

In the training phase, we habituated the baboons to the food boxes, trained prospective specialists to use the apparatus, and selected the male that would be our specialist in the experimental phases to come. During training, the apparatuses only dispensed food directly in front of the lever's position so only the individual training was rewarded. We used shaping techniques to guide the baboons to first approach, then touch the apparatus, then to touch, and fully depress the lever. Any baboon that approached the box could participate in training and receive food rewards regardless of age or sex class.

Field apparatus description

The upper compartment containing the food and electronics was supported by two 10mm clear plastic panels. A steel lever 11cm long protruded from the front side of the box. When the lever was pulled from the full upright position to the lowest position it triggered a servo, which spun, releasing food from the bottom of the upper compartment from where it fell evenly into the four areas created by the crossed base panels.

Model Formulations

1. Approaches to specialist ~ cumulative feeding time*post-day-number*unit-member + precipitation +offset(log(duration present in group)) +
(1+ cumulative feeding time*post-day-number*unit-member + precipitation |subject ID) +
(1+ cumulative feeding time*post-day-number*unit-member + precipitation |specialist ID)

2.Grooming duration of specialist ~ cumulative feeding time*post-day-number + precipitation + (1+ cumulative feeding time*post-day-number + precipitation |subject ID) + (1+ cumulative feeding time*post-day-number + precipitation |specialist ID)

3.Unit female aggressions near specialist ~ cumulative feeding time*post-day-number + precipitation + offset(log(duration present near specialist)) + (1+ cumulative feeding time*post-day-number + precipitation |subject ID) + (1+ cumulative feeding time*post-day-number + precipitation |specialist ID)

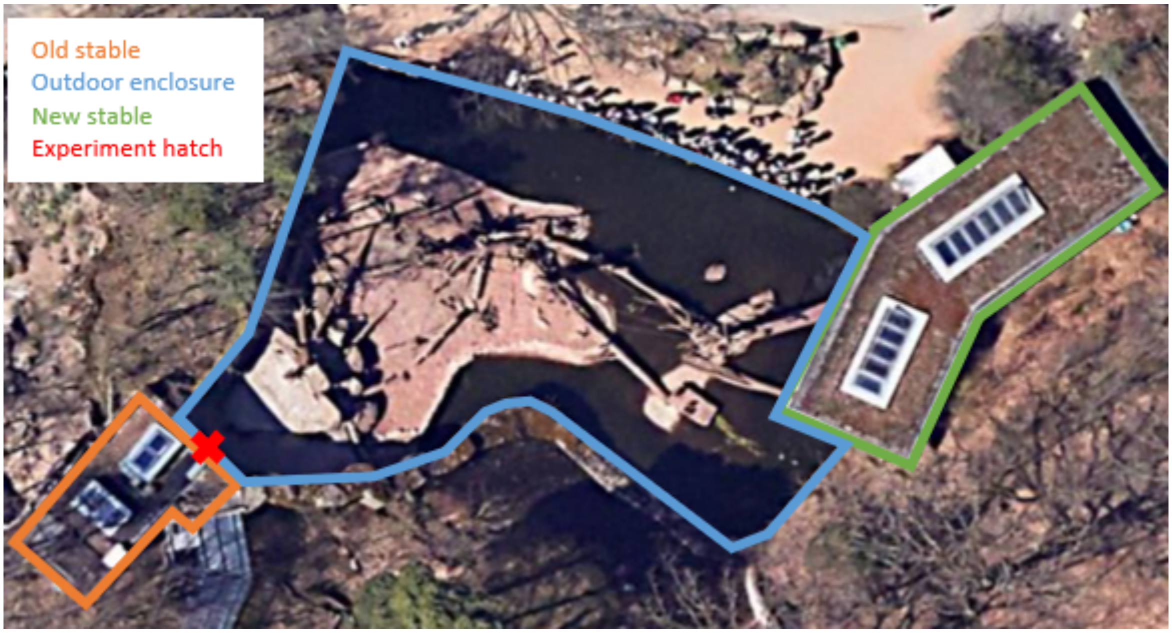


**Fig. S1.**
Satellite view of Guinea baboon housing at Nurnberg Zoo with the three areas and experimental hatch marked (Image from Google Earth).

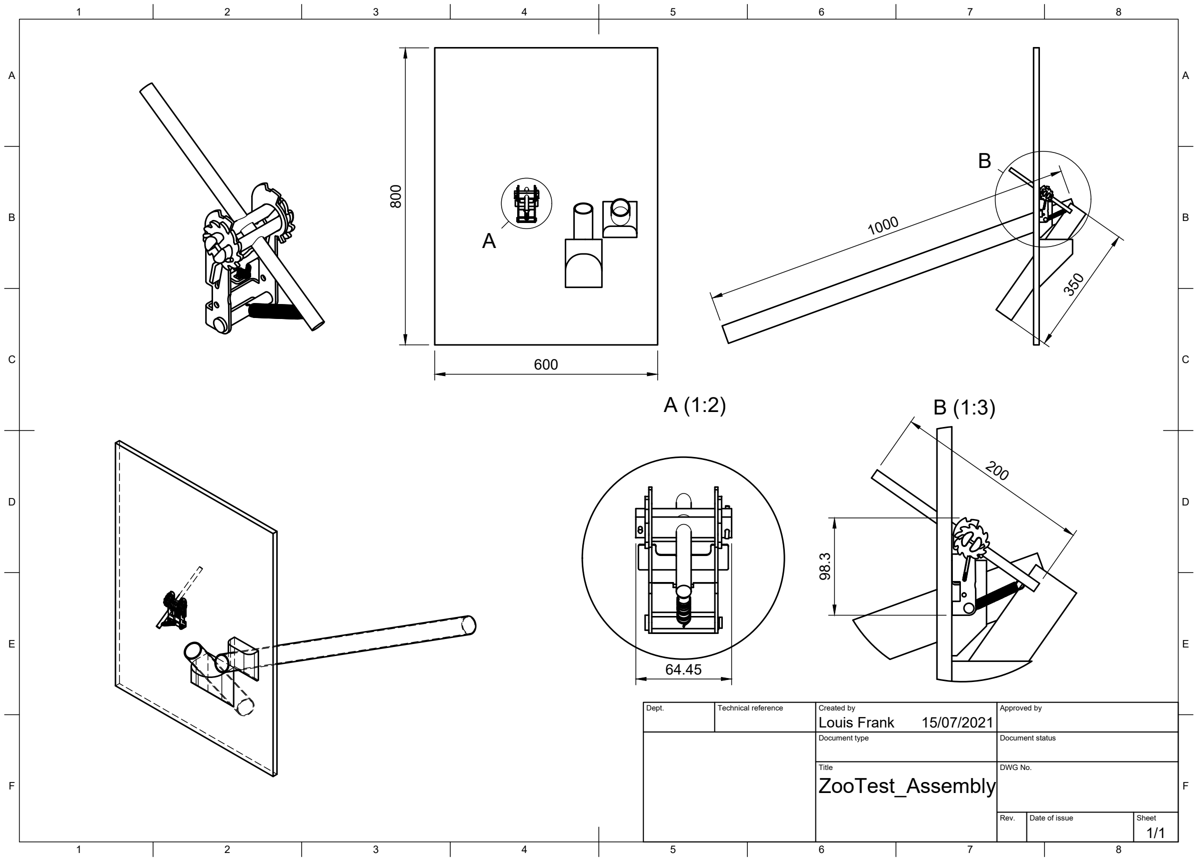


**Fig. S2.**

The experimental apparatus used with the population of Guinea baboons at Nurnberg Zoo. This panel was mounted such that the lever and two pipes were accessible to the baboons. An operator manually dropped peanuts down the pipes as rewards for lever pulls.

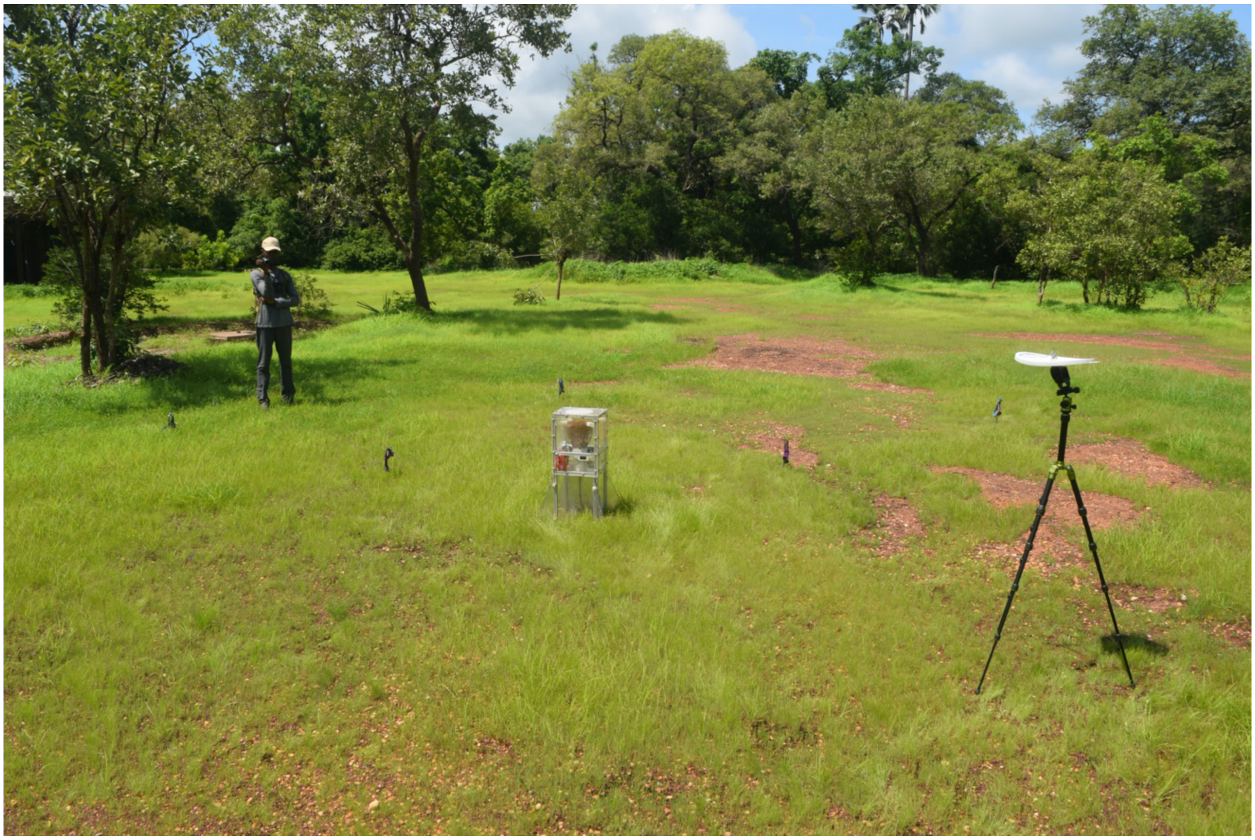

**Fig. S3.**
Experimental set up used with wild baboons at the field site in Senegal.

**Table S1.**

Table showing the unit and party associations of the Guinea baboons within the Tiergarten Nürnberg baboon party October 2021 and the baboons of parties 5 and 6W at our field site in Niokolo Koba National Park in August 2022.

| Full name | ID | Sex | Age category | Gang | Party | Unit |
|---|---|---|---|---|---|---|
| Fefe | FFE | M | Adult | Simenti | 5 | FFE |
| Marleen | MLE | F | Adult | Simenti | 5 | FFE |
| Penny | PNN | F | Adult | Simenti | 5 | FFE |
| Asha | ASA | F | Adult | Simenti | 5 | SPC |
| Fanta | FTA | F | Adult | Simenti | 5 | SPC |
| Kalissi | KLS | F | Adult | Simenti | 5 | SPC |
| Sally | SLY | F | Adult | Simenti | 5 | SPC |
| Spencer | SPC | M | Adult | Simenti | 5 | SPC |
| Amidala | AMD | F | Adult | Simenti | 5 | SPP |
| Anne | ANE | F | Adult | Simenti | 5 | SPP |
| Liselotte | LSL | F | Adult | Simenti | 5 | SPP |
| Quendolin | QND | F | Adult | Simenti | 5 | SPP |
| Sepp | SPP | M | Adult | Simenti | 5 | SPP |
| Jyn | JYN | F | Adult | Simenti | 5 | VDR |
| Rey | REY | F | Adult | Simenti | 5 | VDR |
| Vador | VDR | M | Adult | Simenti | 5 | VDR |
| Abu | ABU | M | Juvenile | Simenti | 5 | |
| Moritz | MRZ | M | Juvenile | Simenti | 5 | |
| Crawford | CRW | M | Adult | Mare | 6W | CRW |
| Effie | EFF | F | Adult | Mare | 6W | CRW |
| Dumbo | DMB | M | Adult | Mare | 6W | |
| Chaplin | CHP | F | Adult | Mare | 6W | LOU |
| Erika | EKA | F | Adult | Mare | 6W | LOU |
| Imogen | IMG | F | Adult | Mare | 6W | LOU |
| Louis | LOU | M | Adult | Mare | 6W | LOU |
| Luna | LUN | F | Adult | Mare | 6W | LOU |
| Xena | XNA | F | Adult | Mare | 6W | LOU |
| Punky | PKY | F | Adult | Mare | 6W | WLD |
| Emilia | EML | F | Adult | Mare | 6W | WLD |
| Ewine | EWN | F | Juvenile | Mare | 6W | WLD |
| Waldo | WLD | M | Adult | Mare | 6W | WLD |
| Charlie | CHR | M | Adult | Mare | 6W | |
| Dita | DIT | M | Juvenile | Mare | 6W | |
| Dr. Rebel | ZDRR | M | Adult | NA | ZOO | ZDRR |
| Luja | ZLUJ | F | Adult | NA | ZOO | ZDRR |

| | | | | | | |
|---|---|---|---|---|---|---|
| Ayla | ZAYL | F | Adult | NA | ZOO | ZDSH |
| Dash | ZDSH | M | Adult | NA | ZOO | ZDSH |
| Dula | ZDUL | F | Adult | NA | ZOO | ZDSH |
| Summer | ZSMM | F | Adult | NA | ZOO | ZDSH |
| Seven | ZSVN | F | Adult | NA | ZOO | ZDSH |
| Grimace | ZGRM | M | Adult | NA | ZOO | ZGRM |
| Marie | ZMAR | F | Adult | NA | ZOO | ZGRM |
| Nada | ZNAD | F | Adult | NA | ZOO | ZGRM |
| Ancient one | ZANC | F | Adult | NA | ZOO | ZHND |
| Handlebar | ZHDB | F | Adult | NA | ZOO | ZHND |
| Handsome | ZHND | M | Adult | NA | ZOO | ZHND |
| Sestra | ZSST | F | Adult | NA | ZOO | ZHND |
| Ela | ZELA | F | Adult | NA | ZOO | ZHNG |
| Hangover | ZHNG | M | Adult | NA | ZOO | ZHNG |
| Smilla | ZSML | F | Adult | NA | ZOO | ZHNG |
| Etti | ZETT | F | Adult | NA | ZOO | ZLCK |
| Lickety | ZLCK | M | Adult | NA | ZOO | ZLCK |
| Louis XVI | ZLSX | M | Adult | NA | ZOO | ZLSX |
| Netta | ZNTT | F | Adult | NA | ZOO | ZLSX |
| Tsau | ZTSA | F | Adult | NA | ZOO | ZLSX |
| Toothless | ZTLS | M | Adult | NA | ZOO | ZTLS |
| Runner | ZRNN | F | Adult | NA | ZOO | ZTLS |
| Four Jane | ZFRJ | F | Adult | NA | ZOO | ZTLS |
| Masked Man | ZMSK | M | Adult | NA | ZOO | |
| Scar Jo | ZSCJ | F | Adult | NA | ZOO | |
| Twig | ZTWG | F | Adult | NA | ZOO | |
| Twain | ZTWN | M | Adult | NA | ZOO | |
| Cloud | ZCLD | M | Adult | NA | ZOO | |
| Thunder | ZTHN | M | Adult | NA | ZOO | |

**Table S2.**
Summary statistics of scans presence and feeding for all males present for at least one box presentation.

| specialist party[1] | subject[1] | party[2] | scans[3] | feeding (hrs)[4] |
|---|---|---|---|---|
| [5] | FFE | [5] | 15 | 0.914 |
| [5] | SPC | [5] | 17 | 0.785 |
| [5] | MKA | [15] | 12 | 0.712 |
| [5] | VDR | [5] | 9 | 0.383 |
| [5] | BLA | [6I] | 8 | 0 |
| [5] | BYE | [9B] | 0 | 0 |
| [5] | CNV | [17] | 1 | 0 |
| [5] | CRN | [17] | 0 | 0 |
| [5] | CSC | [6I] | 2 | 0 |
| [5] | DMB | [6W] | 1 | 0 |
| [5] | IND | [6I] | 4 | 0 |
| [5] | OPA | [17] | 0 | 0 |
| [5] | QNN | [6I] | 3 | 0 |
| [5] | TCO | [13] | 3 | 0 |
| [5] | VDT | [17] | 0 | 0 |
| [5] | XBO | [13] | 9 | 0 |
| [6W] | CHR | [6W] | 113 | 3.461 |
| [6W] | MKA | [15] | 0 | 0.038 |
| [6W] | SPC | [5] | 0 | 0.024 |
| [6W] | SPP | [5] | 0 | 0.011 |
| [6W] | WLD | [6W] | 11 | 0.009 |
| [6W] | BST | [9B] | 3 | 0 |
| [6W] | BYE | [9B] | 0 | 0 |
| [6W] | CRW | [6W] | 4 | 0 |
| [6W] | DMB | [6I] | 12 | 0 |
| [6W] | NYG | [9B] | 2 | 0 |
| [6W] | OSW | [9B] | 7 | 0 |
| [6W] | VDR | [5] | 0 | 0 |
| [6W] | VDT | [13] | 3 | 0 |
| [6W] | VNC | [9B] | 0 | 0 |
| [ZOO] | ZDSH | [ZOO] | 2 | 0.911 |
| [ZOO] | ZLCK | [ZOO] | 7 | 0.783 |
| [ZOO] | ZHNG | [ZOO] | 2 | 0.554 |
| [ZOO] | ZMSK | [ZOO] | 0 | 0.304 |
| [ZOO] | ZDRR | [ZOO] | 3 | 0.232 |

| | | | | |
|---|---|---|---|---|
| [ZOO] | ZTHN | [ZOO] | 17 | 0.130 |
| [ZOO] | ZLSX | [ZOO] | 0 | 0.036 |
| [ZOO] | ZGRM | [ZOO] | 3 | 0.024 |
| [ZOO] | ZTLS | [ZOO] | 4 | 0.008 |
| [ZOO] | ZTWN | [ZOO] | 8 | 0.007 |
| [ZOO] | ZCLD | [ZOO] | 1 | 0.001 |

[1] “specialist party[1]” indicates the party of the specialist in which they were present for scans and ate from the box.

[2] “subject” to their 3-letter code, “party” refers to the party of the individual,

[3] “scans” is the number of 5 m-proximity scans taken of the specialist in which the individual was present. Total scans taken in each party:[6W] = 817, [5] = 589, [ZOO] =165.

**Movies S1-S3.**

The following link is to a playlist of clips from food box presentations with the Guinea baboon three parties.
https://www.youtube.com/playlist?list=PLwD_i507VU-p-ZW1MBQVJwYJ3hJGsLeAw
